# Heparin as an Anti-Inflammatory Agent

**DOI:** 10.1101/2020.07.29.223859

**Authors:** Leandar Litov, Peicho Petkov, Miroslav Rangelov, Nevena Ilieva, Elena Lilkova, Nadezhda Todorova, Elena Krachmarova, Kristina Malinova, Anastas Gospodinov, Rossitsa Hristova, Ivan Ivanov, Genoveva Nacheva

## Abstract

Timely control of the cytokine release syndrome (CRS) at the severe stage of COVID-19 is key to improving the treatment success and reducing the mortality rate. The inhibition of the activity of the two key cytokines, IFNγ and IL-6, can significantly reduce or even reverse the development of the cytokine storm. The objective of our investigations is to reveal the anti-inflammatory potential of heparin for prevention and suppression of the development of CRS in acute COVID-19 patients.

The effect of low-molecular-weight heparin (LMWH) on IFNγ signalling inside the stimulated WISH cells was investigated by measuring its antiproliferative activity and the translocation of phosphorylated STAT1 in the nucleus. The mechanism of heparin binding to IFNγ and IL-6 and therefore inhibition of their activity was studied by means of extensive molecular-dynamics simulations. We find that LMWH binds with high affinity to IFNγ and is able to inhibit fully the interaction with its cellular receptor. It also influences the biological activity of IL-6 by binding to either IL-6 or IL-6/IL-6Rα thus preventing the formation of the IL-6/IL-6Rα/gp130 signaling complex. Our conclusion is that heparin is a potent anti-inflammatory agent that can be used in acute inflammatory conditions, due to its potential to inhibit both IFN γ and IL-6 signalling pathways. Based on our results and available clinical observations, we suggest the administration of LMWH to COVID-19 patients in the initial stages of the acute phase. The beginning of the treatment and the dosage should be based on a careful follow-up of the platelet count and the D-dimer, IL-6, IFN, T-cells, and B-cells levels.

## 1 Introduction

COVID-19, the disease caused by the new coronavirus SARS-CoV-2 [1], accounts so far for over 16 million cases worldwide and more than 650000 deaths. The majority of COVID-19 patients have no or only mild symptoms. However, about 15% of them develop severe symptoms and many require intensive care treatment. SARS-CoV-2 infection can be roughly divided into three stages: stage I, an asymptomatic incubation period with or without detectable virus; stage II, a non-severe symptomatic period; stage III, severe respiratory symptomatic stage with a high viral load [2]. Until now there are no specific drugs approved as an effective means for fighting COVID-19.

Clinically, the immune responses induced by SARS-CoV-2 infection have two phases. During the incubation and non-severe stages, a specific innate immune response is required to eliminate the virus and to prevent disease progression to severe stages. Later, the adaptive immune response is activated to completely clear the remaining viruses. The average incubation period of COVID-19 is estimated to be between 6 and 13 days [3] – longer than the one for influenza, suggesting that SARS-CoV-2 may have developed mechanisms to counteract the immune response, such as delaying the type I interferon response and thus inhibiting innate immune signalling [4].

Severe COVID-19 disease manifested through fever and pneumonia, leading to acute respiratory distress syndrome (ARDS) and accompanied by a cytokine release syndrome (CRS), has been described in up to 20% of the cases [5]. The cytokine storm and the subsequent ARDS result from the combined action of many immunoactive molecules. Interferons, interleukins, and chemokines are the main components involved in the development of the cytokine storm [6,7]. Already the first investigated patients in China showed leucopenia, lymphopenia, and high values of prothrombin time and D-dimer level during the severe phase. The plasma levels of IL-1B, IL-1RA, IL-6, IL-7, IL-8, IL-9, IL-10, basic FGF, GCSF (CSF3), GMCSF (CSF2), IFNγ, IP-10 (CXCL10), MCP-1 (CCL2), MIP-1A (CCL3), MIP-1B (CCL4), PDGF, TNF-α, and VEGF concentrations were higher in both intensive-care-unit (ICU) patients and non-ICU patients than in healthy adults [5]. Earlier studies have shown that increased serum levels of proinflammatory cytokines (e.g., IL-1B, IL-6, IL-12, IFNγ, IP-10, and MCP-1) were associated with pulmonary inflammation and extensive lung damage in SARS-CoV and MERS-CoV patients [8,9].

Timely control of the cytokine storm in its early stage is the key to improving the treatment success rate and reducing the mortality rate in patients with COVID-19. Recently, a number of specific anti-cytokine approaches have been proposed for the treatment of the CRS, based on drugs targeting IL-1, IL-6, IL-18, and IFNγ [10]. High IL-6 levels are strongly correlated with some of the prominent symptoms of cytokine storm, such as vascular leakage, activation of the complement and coagulation cascades, inducing disseminated intravascular coagulation. The inhibition of the activity of the two key cytokines, IFNγ and IL-6, can significantly reduce or even reverse the development of the cytokine storm. One possible candidate able to bind to both cytokines is low-molecular-weight heparin (LMWH).

Interferon-gamma (IFNγ) is a pleiotropic cytokine. The mature form of human IFNγ is a non-covalent homodimer, each of the monomers consisting of 143 amino acids (aa), 62% of which are arranged in 6 α-helices (A to F), associated with short unstructured regions and highly flexible C-terminal end (21 aa). Upon binding to its receptor, IFN**γ** activates the JAK/STAT1α signal transduction pathway. STAT1 is phosphorylated, subsequently dimerises and then translocates into the nucleus to initiate the transcription of target genes. The 3D structure of the complex of IFNγ with its receptor, which consists of two non-covalently associated chains, IFNGR1 and IFNGR2, is resolved and the binding sites are mapped on the IFNγ molecule [11,12]. They are located in the loop between helices A and B (residues 18–26), His111 and a short putative area (residues 128–131) in the flexible C-terminal domain.

Interleukin 6 (IL-6) is a member of the IL-6 cytokine family, secreted by many cell types upon stimulation during infection, inflammation or cancer [13]. IL-6 is important for regulating B cell and T cell responses and for coordinating the activity of the innate and the adaptive immune systems. IL-6 is a 26 kDa, 212 residues bundle cytokine shaped in 4 α-helical parts with two extensive loop regions situated between helices A and B and helices C and D [14]. It binds to the IL-6 receptor (IL-6R, also known as IL-6R subunit-α or IL-6Rα), an 80 kDa protein devoid of signalling capacity. This complex binds to a second membrane protein, glycoprotein 130 (gp130, also known as IL-6R subunit-ß and acting as a signalling receptor for the whole IL-6 family), which dimerises and initiates intracellular signalling [15]. gp130 is ubiquitously expressed on the cell surface, however IL-6Rα is found only on a few cell types, such as hepatocytes, some leukocytes and epithelial cells [16]. IL-6 exhibits affinity only for IL-6Rα but not for gp130. The complex IL-6/IL-6Rα/gp130 homodimerises to form a hexamer that is a prerequisite for the initiation of the IL-6 intracellular signaling.

Heparin is a naturally occurring polysaccharide, a member of the glycosaminoglycan (GAG) family. It is widely used in the medical practice as an anticoagulant, with common mild side effects but also some more severe ones, among them the heparin-induced thrombocytopenia [17]. Heparin is a linear highly sulfated polysaccharide chain, composed of repeating disaccharide units of 1,4 linked α-L-iduronic or β-D-glucuronic acid (D-GlcA), and α-D-glucosamine (D-GlcN). The predominant substitution pattern comprises 2-O-sulfation of the iduronate residues and N- and 6-O-sulfation of the glucosamine residues [18,19]. This structure defines one of the most prominent characteristics of the molecule – a high density negative electric charge. This is the main reason for the high biological activity of the heparin. It binds more than 700 proteins. While natural heparin (also known as UFH or unfractionated heparin) consists of chains of varying molecular weight, from 5 kDa to over 40 kDa, its fractionated form, the low molecular weight heparin (LMWH), consists of more uniformly weighted chains – at least 60% of them being below 8 kDa of weight, and is preferred in the clinical practice because of its more predictable pharmacokinetics and milder side effects.

In a biological context, heparin action is more closely linked to defense against exogenous pathogens [20] and responses following tissue damage. Heparin derivatives were shown to be capable of acting at several points in the inflammatory process [21]. Heparin alters cytokine levels [22] and one of its biological roles may be to dampen the effects of the sudden release of large numbers of cytokines following infection or sudden trauma.

The objective of our investigations is to reveal the anti-inflammatory potential of heparin for prevention and suppression of the development of cytokine storm in acute COVID-19 patients. In particular, we investigate its ability to bind to IFNγ and IL-6 cytokines and inhibit their signaling pathways, thus opening the door for modulation of the development of cytokine storm.

## 2 Materials and methods

### 2.1 Explicit solvent MD simulations

The MD simulations in this study were performed with the GROMACS software package [23– 25]. If not specified otherwise, the MD protocol consisted of a force field parameterization, structure-in-vacuum energy minimization, solvation, charge neutralization through Na^+^ and Cl^-^ ions at 0.15 M concentration, energy minimization, MD run with position restraints on heavy solute atoms (PR run) and production MD runs in the NPT ensemble. The temperature was kept constant by applying Berendsen [26] and V-rescale [27] thermostats during PR and production run, respectively. Berendsen barostat was used in equilibration MD runs and Parrinello-Rahman barostat – for the production MD runs [28]. Long-range electrostatic interactions were treated by applying the Particle Mesh Ewald algorithm [29] with 1.2 nm cut-off radius for short-range interactions. Switch function was used for Van der Waals interactions calculations with 1.2 nm cut-off radius and 1 nm switching distance. We used the leap-frog integration algorithm with 2 fs time step, the covalent bonds connecting hydrogen atoms with the solute heavy atoms were kept with fixed length by LINCS algorithm [30] and RATTLE algorithm [31] was used to keep the solvent molecules rigid.

### 2.2 Heparin hexasaccharide structure

Based on data from literature [32–34] the following hexasaccharide sequence was chosen as a model molecule for a general LMWH: GlcNAc(6S) (1→4) GlcA (1→4) GlcNS(6S) (1→4) IdoA(2S) (1→4) GlcNS (1→4) GlcA(2S). This oligosaccharide chain has a net charge of -9e. The Glycan Reader & Modeler module [35] of the CHARMM-GUI server [36] was used for generation of a three-dimensional structure, corresponding to the chosen carbohydrate sequence, as well as a topology using the latest version of the CHARMM36 carbohydrate force field [37]. The topology was converted to a GROMACS-compatible topology using the parmed module of Ambertools 16 [38].

### 2.3 Human interferon gamma

The 3D model of reconstructed and folded full-length IFNγ [39,40] was used as the initial structure for the IFNγ-associated simulations.

### 2.4 Human interleukin 6

The crystal structure of human interleukin-6 with PDB ID 1 ALU [41] was taken from [42]. The missing residues, ^52^SerSerLysGluAlaLeuAlaGluAsn^60^, were added using the loop-modeling interface [43] to MODELLER [44] of UCSF Chimera [45]. A 10 ns production MD run was performed to prepare the initial structure of the IL-6 molecule for all subsequent IL-6 related studies.

### 2.5 IL-6/IL-6Rα/gp130 complex

The crystal structure of human IL-6/IL-6Rα/gp130 complex with PDB ID 1P9M [46] was taken from [42]. The 3D structure for interaction analysis was prepared by a 10 ns MD simulation as described above.

### 2.6 Heparin-protein complexes simulations

To study the interaction of IFNγ with LMWH, four hexasaccharides were placed near the two C-termini of the homodimer. The minimal distance between the carbohydrates and the protein was 1.6 nm. The MOE software package was employed to prepare the initial structures of the complexes of IL-6 and heparin, and IL-6/IL6Rα and heparin. The heparin molecules were docked in IL-6 and IL-6/IL6Rα complex based in proximity to IL-6 binding sites I and II [46].

### 2.7 Complex-interactions Analysis

The evaluation of the interactions in the investigated molecular complexes was performed with the MOE software suit. To this end, the MD-generated structures were further optimised in the MMFF94 force field to allow for a hydrophobicity profiling of the respective SAS. The interactions between IL-6 or IL-6/IL6Rα and the heparin molecules were analysed and visualised with the ligand interactions functionality of MOE [47,48].

### 2.8 Cell culture

WISH cell line (ATCC® CCL-25™) was propagated in Eagle’s Minimum Essential Medium (EMEM, ATCC® 30-2003™) supplemented with 10% fetal bovine serum (Gibco™) and penicillin-streptomycin (10000 U/ml, Gibco™). Cells were cultured in 25-cm^2^ flasks (Thermo Scientific™ Nunc™) at 37°C incubator with 5% CO2.

### 2.9 Inhibitory antiproliferative assay

Confluent WISH cell monolayers were detached by trypsin-EDTA treatment (Sigma) and were seeded in a 96 well plate (Corning®) at a density of 1 × 10^6^ cells/ml. Cells were treated with: i) 100 ng/ml IFNγ purified as described in [49], ii) IFNγ (100 ng/ml) supplemented with Fraxiparine (GlaxoSmithKline Pharmaceuticals S.A.) in concentrations varying from 1.5 to 150 anti-Xa IU/ml pre-incubated for 1h at room temperature and iii) Fraxiparine in the same concentrations as in ii). Further the antiproliferative activity of IFNγ was measured by a modified kynurenine bioassay as described earlier [50].

### 2.10 Cellular localisation of phosphorylated STAT1

In order to determine the localisation of the phosphorylated STAT1, WISH cells were grown on coverslips, washed twice with 1 × PBS and treated for 15 min with 100 ng/ml IFNγ alone or IFNγ (100 ng/ml) pre-incubated for 1h at room temperature with Fraxiparine in concentrations varying from 7.5 to 150 anti-Xa IU/ml. As a control, the cells were separately treated with the highest concentration of Fraxiparine (150 anti-Xa IU/ml). Following the incubation, the cells were first fixed with 3.7% formaldehyde in 1 × PBS for 7 min at room temperature followed by washing with 1 × PBS. Next, the cells were fixed with methanol for 5 min at – 20°C. After the fixation, the cells were permeabilised with 0.5% Triton X-100 in PBS for 5 min, washed with 1 × PBS and blocked for 1 h at room temperature with blocking buffer (5% BSA, 0.1% Tween 20 in 1 × PBS). The cells where incubated overnight at 4°C with rabbit anti-phospho-STAT1 (Tyr701) primary antibody (Cell Signaling Technology, 9167S) diluted 1:100 in blocking buffer, followed by washing 5 times with 0.1% Tween 20 in 1 × PBS. Cells were then stained with secondary antibody Alexa Fluor® 594 goat anti-rabbit IgG in blocking buffer at dilution 1:500 for 1 h at room temperature in the dark. Following the incubation, the coverslips were washed 4 times with 0.1% Tween 20 in 1 × PBS. The nuclei were counterstained with 0.5 μg/ml DAPI (Sigma-Aldrich) for 2 min at room temperature. The coverslips were washed with distilled water and mounted on glass slides with ProLong™ Gold Antifade mounting media (Invitrogen). Images were analysed by CellProfiler software [51].

## 3 Results

### 3.1 Effect of heparin on the activity of IFNγ

In this study the antiproliferative activity of IFNγ was used as a basis to study the effect of LMWH. It was measured by a modified kynurenine bioassay [50] run with a referent recombinant IFNγ having specific biological activity of 10^6^ IU/mg [49]. Preformed monolayers of WISH cells were treated with 100 ng/ml (100 IU/ml) IFNγ alone or pre-incubated with various concentrations of LMWH. The antiproliferative activity of IFNγ in this assay is due to the induction of indoleamine-2,3-dioxygenase (IDO), which is the first and rate limiting enzyme in the tryptophan catabolism, catalysing oxidative cleavage of tryptophan to N-formylkynurenine. Following a hydrolysis step, the latter is transformed into kynurenine, which results with Ehrlich’s reagent in a yellow-coloured compound absorbing at 490 nm. As seen in Fig. 1, LMWH inhibited the antiproliferative activity of IFNγ in a concentration-dependent manner with concentration of about 35 IU/ml reducing it to 50% of the control level. The addition of LMWH at 150 IU/ml was not toxic to the cells but abolished the induction of IDO up to 80%.

**Fig. 1.**
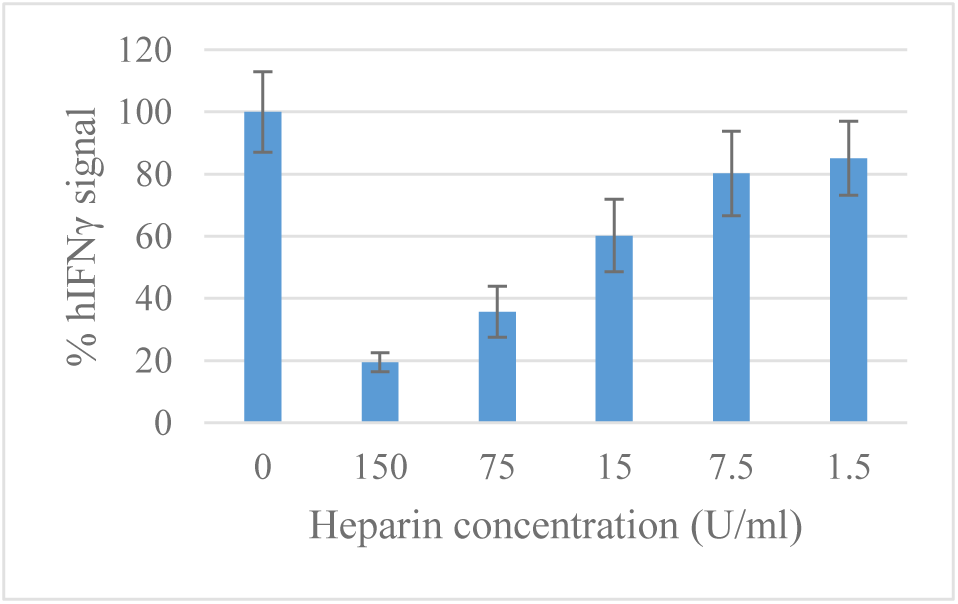
Inhibitory effect of LMWH on the production of IDO induced by 100 IU/ml recombinant IFNγ. After subtraction of the blank value, the absorbance obtained from cells treated with a mixture of IFNγ and LMWH is related to that obtained from cells treated with IFNγ only, taken as 100%. The bars represent the statistical errors.

In order to further investigate the effect of LMWH on IFNγ signalling inside the stimulated cells we studied the translocation of phosphorylated STAT1 in the nucleus using specific antibodies. The endogenous phosphor-STAT1 (Tyr^701^) was visualised by immuno-fluorescence staining of fixed WISH cells after 15 min induction with 100 ng/ml (100 IU/ml) IFNγ alone or in the presence of various concentrations of LMWH (Fig. 2). The obtained results once again confirmed that LMWH abolishes the IFNγ signalling, showing full inhibition of the phosphorylation of STAT1 and therefore its translocation into the nucleus at 150 U/ml.

**Fig. 2.**
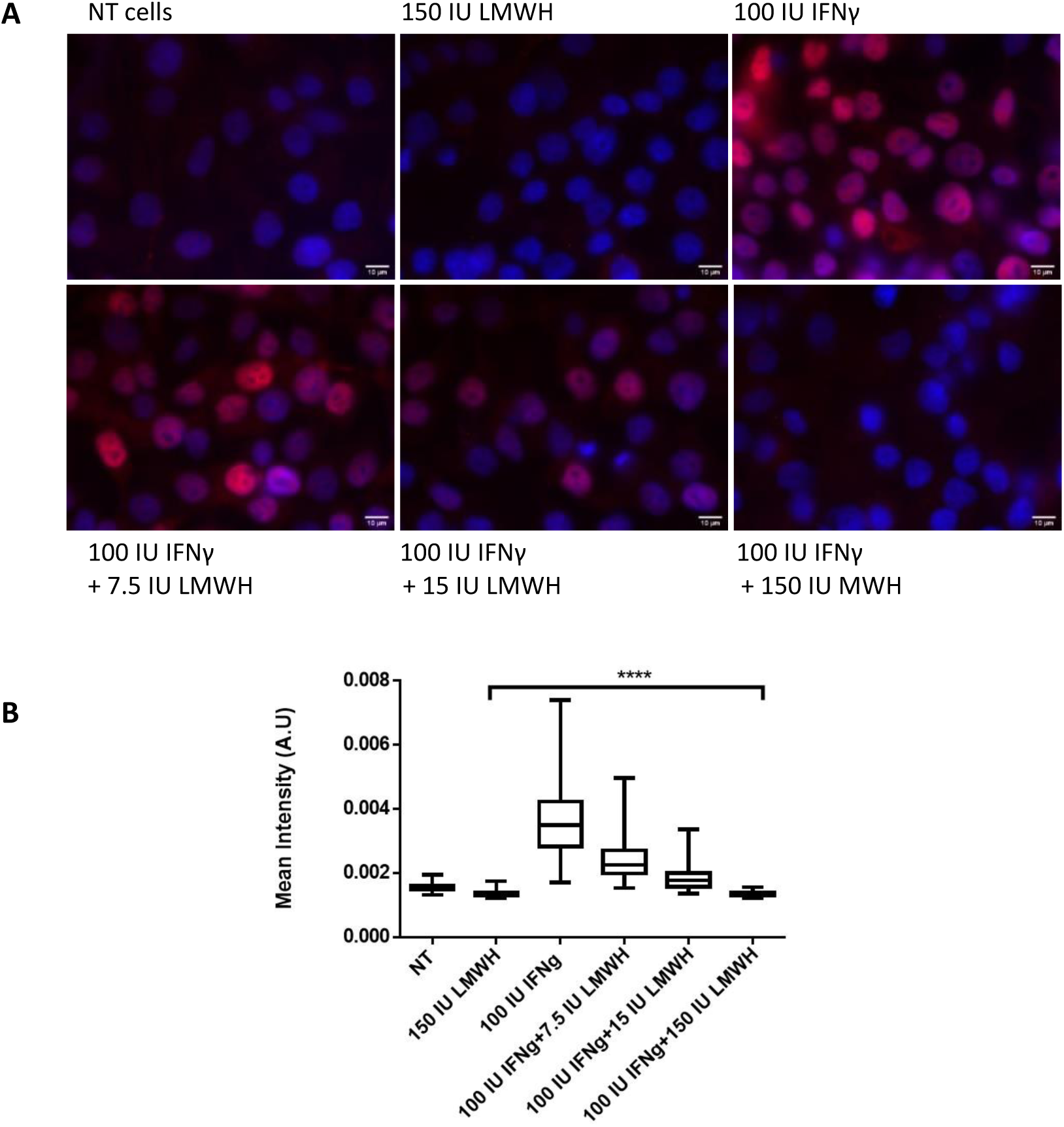
Translocation of the phosphorylated STAT1 into the cell nucleus. (A) WISH cells were treated for 15 min with 100 ng/ml IFNγ or IFNγ (100 ng/ml), pre-incubated with LMWH at concentrations 7.5, 15 and 150 anti-Xa IU/ml. Non-treated cells (NT cells) and cells treated with 150 anti-Xa IU/ml LMWH (150 IU LMWH) were used as controls. After treatment cells were collected, fixed, and stained with an antibody against pSTAT1 (Tyr701). The nuclei were stained with DAPI. Representative images are shown, scale bar 10 µm. (B) Intensity of pSTAT1 (Tyr701) staining of individual nuclei from (A). Data are averaged from 3 independent experiments; ****P < 0.0001 (two-tailed unpaired Student’s t-test).

The obtained results indisputably show that LMWH abolishes the biological activity of IFNγ. In order to shed light on the mechanism of action of LMWH we performed MD simulation of the interaction of heparin hexasaccharides with the IFNγ molecule.

### 3.2 Simulation of heparin hexasaccharides’ interaction with IFNγ

IFNγ is a basic cytokine, with the full-length homodimer having a net charge of +18e. The positive charge is concentrated mainly in the domains D1 (^125^LysThrGlyLysArgLysArg^131^, charge +5e) and D2 (^137^ArgGlyArgArg^140^, charge +3e) of the C-tails. LMWH sugars are highly negatively charged, each chain having a net charge of -9e. Expectedly, the interaction between IFNγ and the hexasaccharides must be dominated by a very intensive electrostatic attraction.

The MD simulation of 500 ns (the initial and the final conformations shown in Fig. 3) confirmed this expectation. The first carbohydrate attaches to the cytokine in just 1 ns. Another nanosecond later, three of the four saccharides have formed at least one contact with the cytokine. The number of contacts between the protein and the sugars is shown in Fig. 4. Within the next 35- 40 ns, the carbohydrate chains move closer and closer to the protein molecule and form more and more contacts, until all three sugars lie down on the protein surface and the number of contacts curve plateaus. This is reflected by the solvent accessible surface area of the carbohydrates, shown in Fig. S1 in the Supplementary Material. The fourth LMWH chain binds to IFNγ after 204 ns. It should be noted, that by that time the complex formed between IFNγ and the first three hexasaccharides already has a net negative charge. As with the first three carbohydrates, after the initial binding this fourth sugar quickly adopts a conformation, covering the cytokine surface, which saturates the number of contacts.

**Fig. 3.**
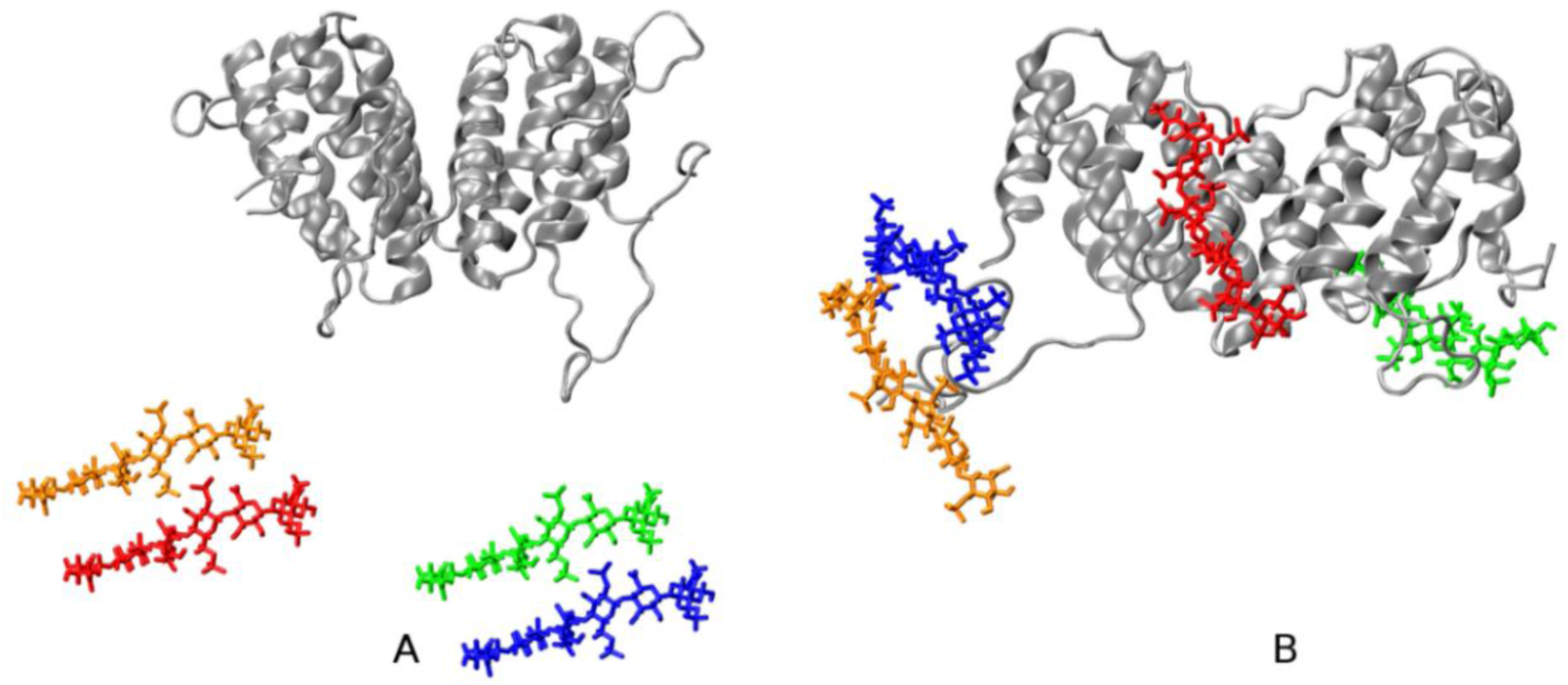
Initial (A) and final (B) configurations of the IFNγ–LMWH hexasaccharides binding simulation. The protein is shown in gray ribbon, the hexasaccharide chains are coloured as follows: dp6_1 – green, dp6_2 –orange, dp6_3 – blue, and dp6_4 – red.

**Fig. 4.**
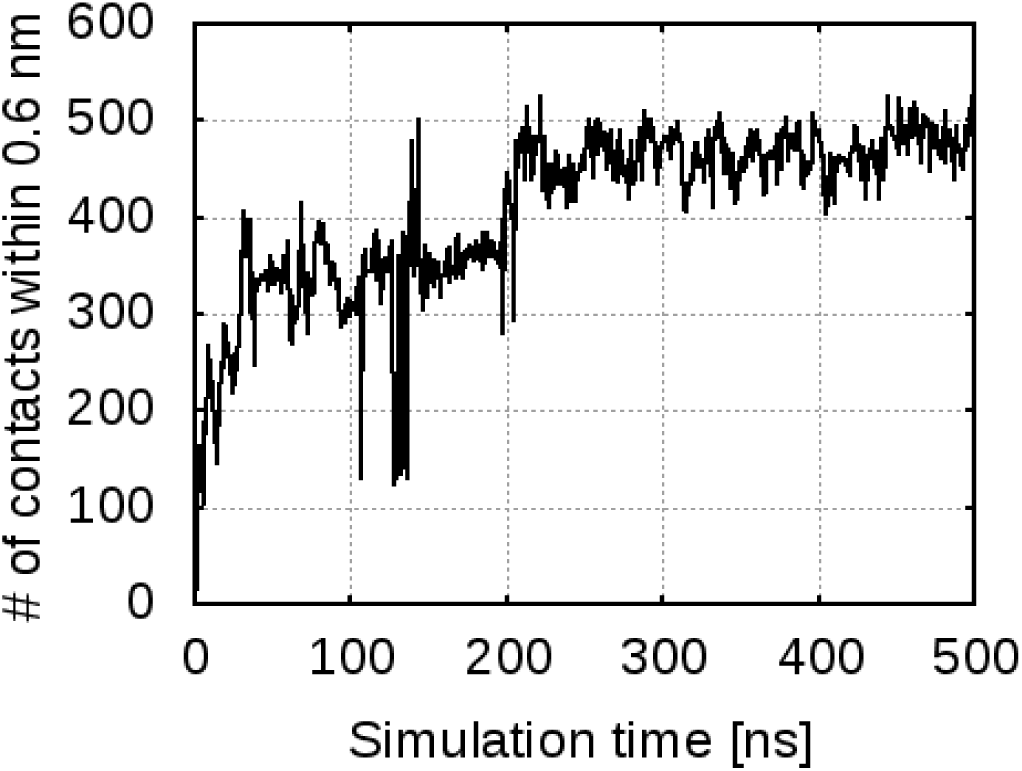
Pair contacts between IFNγ and the hexasaccharides. Number of contacts as a function of the simulation time between any pair of atoms of IFNγ and any of the four hexasaccharides within 0.6 nm.

The IFNγ/LMWH saccharides complexes are extremely stable. Indeed, once a few contacts between a sugar chain and the protein are formed, the sugars do not dissociate from the complex within the simulated time. These complexes are stabilised through a fairly high number of hydrogen bonds. On average in the last 250ns of the simulation, dp6_1 forms 9 +/- 2 H-bonds, dp6_2 – 10+/-2, dp6_3 – 12+/-2 and dp6_4 – 4+/-1. The hydrogen bonds between each hexasaccharide and the IFNγ homodimer are shown in Fig. S2 in the Supplementary Material.

As expected, the LMWH chains interact with the positively charged parts of the IFNγ molecule. The cytokine has three basic clusters – one in the globular part in helix E, ^86^LysLysLysArg^89^, and two in the flexible C-terminal tail, the D1 and D2 regions described above. These are all located in the bottom part of the IFNγ molecule. An electrostatic potential surface of the input protein configuration, showing the surface distribution of charges, is depicted in Fig. S3 of the Supplementary Material.

Fig. 5 shows a contact map of the close interactions of the four hexasaccharides and the two IFNγ monomers. As evident on the map, the sugars bind to exactly these three domains. Dp6_1 binds to the D1 domain in the C-terminus of chain B of the cytokine. Later this C-terminus swings and approaches the globule, which allows dp6_1 to form a few contacts with helix E of chain A. Dp6_2 and dp6_3 both attach to the C-terminus of chain A. Dp6_2 interacts with both the D1 and D2 domains. Dp6_3 binds on the other side of the D1 domain, which places it in close proximity to helix A in chain B, where it forms contacts with the positively charged N-terminal NH3 group of Gln1, Asp2, Lys6 and Lys12-Lys13. The fourth hexasaccharide, dp6_4 binds to the basic cluster in helix E of chain B. As it attaches to the surface of the IFNγ globule, it also forms some loose contacts with the sequence Lys34AsnTrpLys38 at the end of helix B in chain A.

**Fig. 5.**
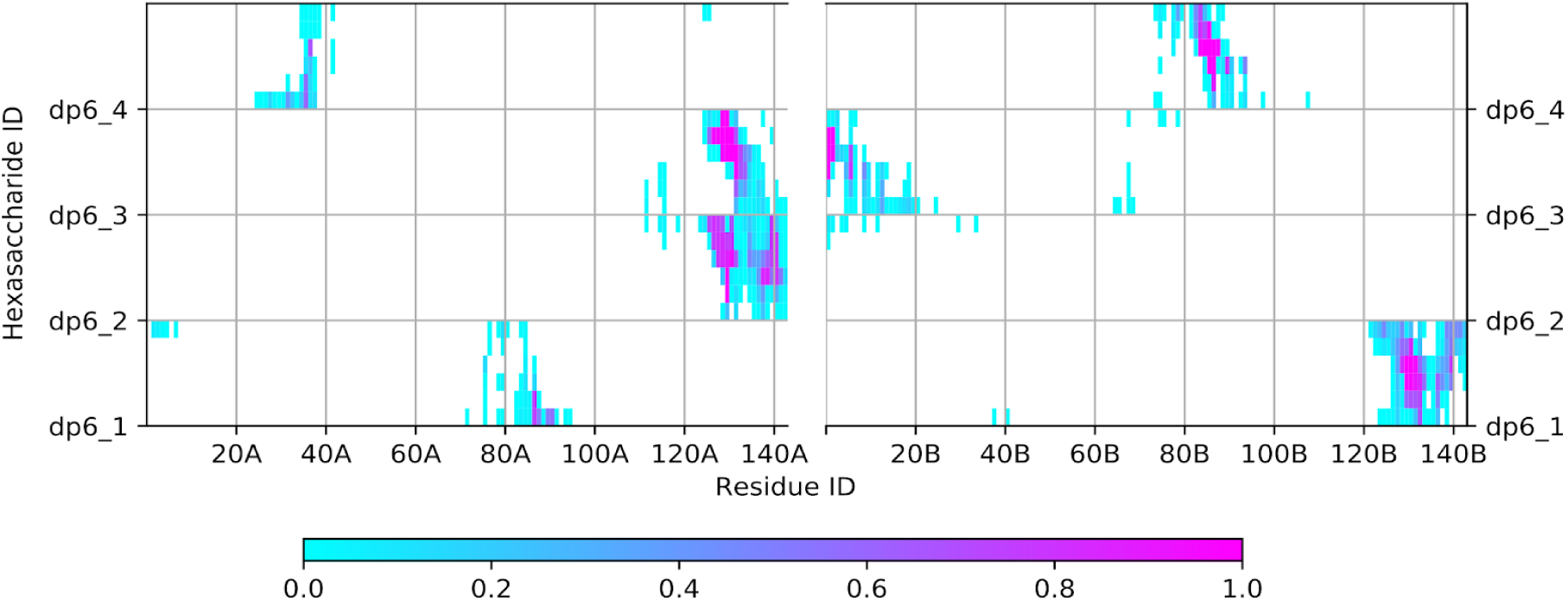
Contact map of the IFNγ /hexasaccharides complex. Contacts within 0.6 nm formed between each of the four hexasaccharides and the two monomers of IFNγ. Contact occupancy within the last 250 ns of the simulation ranges from 0 (cyan) to 1 (purple).

Our simulation shows that LMWH binds to the C-termini of IFNγ with high affinity, forming very stable complexes due to the strong electrostatic attraction. After binding of two or more LMWH hexasaccharides the complex changes its net charge from positive to negative. This impedes further interaction of the cytokine with the extracellular part of the IFNGR1 (also negatively charged) which is the first necessary step in the IFNγ transduction pathway. The *in silico* simulation data explains the obtained experimental results. These findings put forward heparin as an effective inhibitor of IFNγ and, as such, a good candidate for influencing the development of the cytokine storm.

### 3.3 LMWH binding to the IL-6 and IL-6/IL-6Rα complex

As a prerequisite for the dynamics studies, all binding sites in the triple complex IL-6/IL-6Rα/gp130 were scrutinised. The complex structure prepared as described in Sec. 2.6 is shown in Fig. 6. Three binding sites were identified in IL-6 (for details see [52]: Site I, comprising residues Arg30, Leu33, Lys54 (helix A) and Leu178, Arg179, Arg182, Gln183 (helix D), involved in the interaction of IL-6 with IL-6Rα receptor; Site II, comprising residues Arg24, Gln28, Tyr31, Asp34 (helix A) and Glu110, Met117, Val121, Gln124, Phe125 (helix C), involved in the binding of IL-6 to gp130 receptor; Site III, involved in the formation of the hexamer complex.

**Fig. 6.**
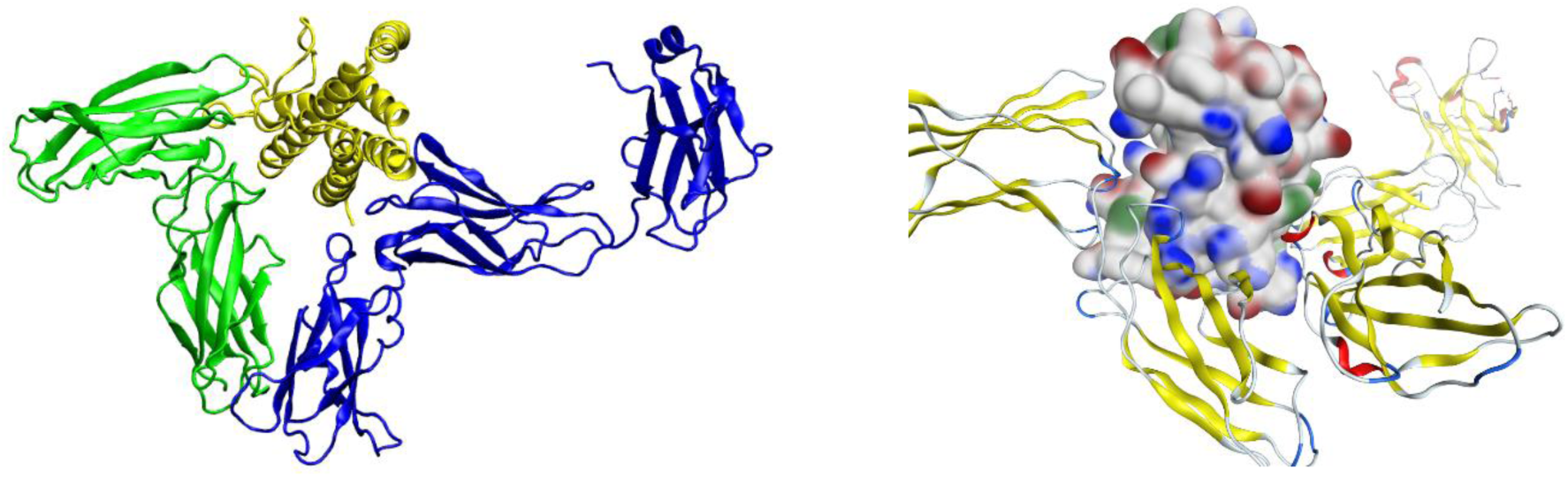
The IL6/IL6Rα/gp130 complex. (A) IL-6 in yellow, IL-6Rα in green, gp130 in blue; (B) IL-6 molecule presented with its SAS (hydrophobic regions in green, positively charged regions in blue, negatively charged regions in red), IL-6Rα and gp130 molecules (in ribbon representation) are to the left, resp. to the right of the IL-6 molecule.

A 250 ns simulation of IL-6/LMWH complex was performed with the initial conformation chosen with account for the identified binding sites, specifically targeting the positively charged amino acid residues of IL-6, involved in purely polar interactions between IL-6 and IL-6Rα (Fig. 7a). A stable conformation was achieved, with heparin binding at Leu19 and Arg30 from helix A and Lys171, Arg179 and Arg182 belonging to helix D. All these interactions are electrostatic in nature, determined by the strong attraction between the positively charged residues of IL-6 and the negatively charged heparin (Fig. 7b). In particular, in this interaction the key residues from site I Arg30, Arg179 and Arg182 are engaged. Actually the heparin molecule is bound at IL-6 helix A and helix D, blocking the IL-6 binding sites to the IL-6Rα thus inhibiting the IL-6 interaction with its receptor.

**Fig. 7.**
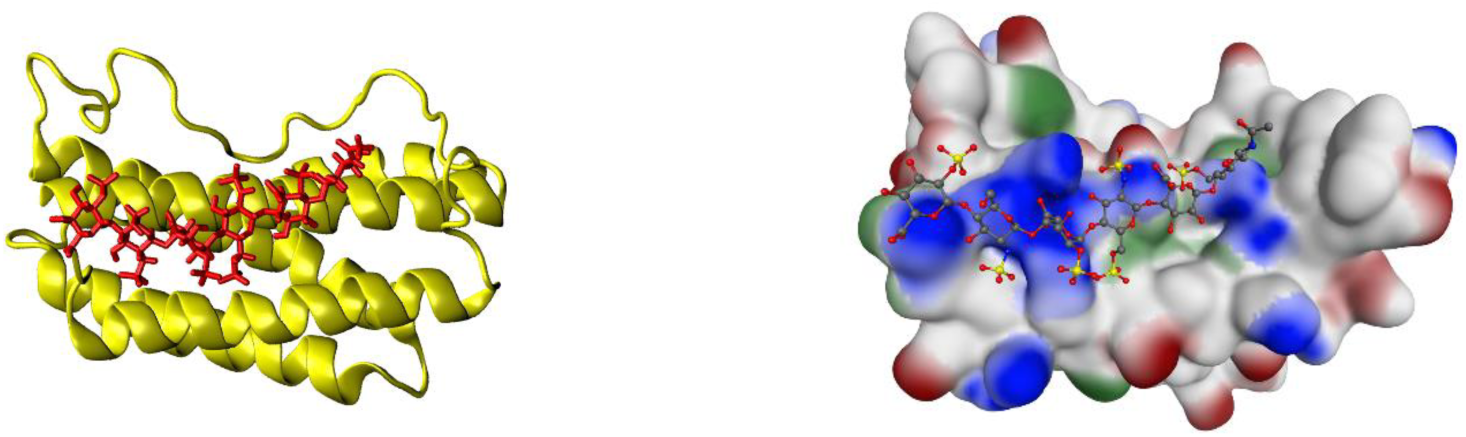
The IL6/LMWH complex. (A) IL-6 in yellow, heparin in red; (B) IL-6 molecule presented with its SAS (hydrophobic regions in green, positively charged regions in blue, and negatively charged regions in red), heparin coloured by atom type.

In studying the possibility of heparin hindering through its binding to the IL-6/IL-6Rα complex the subsequent binding of the latter to gp130, two putative positions were identified. In the first one, heparin is strongly bound to residues Lys40, Lys41, and Arg168 of IL-6, which are not among the direct participants in the complex formation. Thus, heparin impact analysis in this conformation requires full-complex large-scale simulations. In the second position heparin binds to Arg30 of IL-6 helix A and Lys252 of IL-6Rα and comes in close proximity to Tyr31 of IL-6 helix A. This position is of particular interest because, based on mutation studies [53], Tyr31 is considered to be of key importance in the biological activity of IL-6. However, a 250 ns simulation revealed this position as less stable. Its stabilization was achieved by adding two Mg^2+^ ions in another 250 ns simulation, the final conformation of which is depicted in Fig. 8 (left panel). This result is not surprising, taking into consideration the findings about the Ca^2+^-dependent action of heparin outside of the anticoagulant context (see, e.g., [32]). With the gp130 receptor superimposed onto the simulated system (Fig. 8, right panel), we realise the significant impact potential of LMWH on the formation of the IL-6/IL-6Rα/gp130 complex. Not only does heparin bind to Arg30, the immediate neighbour of the key residue Tyr31, but it is also positioned in front of helices A and C of IL-6, thus covering the Tyr31-part of IL-6 binding site II and preventing helix C of IL-6 from getting close enough to gp130 to form a complex. As a consequence, IL-6 and its two receptor parts fail to form a biologically active complex, meaning that the involvement of LMWH interrupts the IL-6 signalling pathway.

**Fig. 8.**
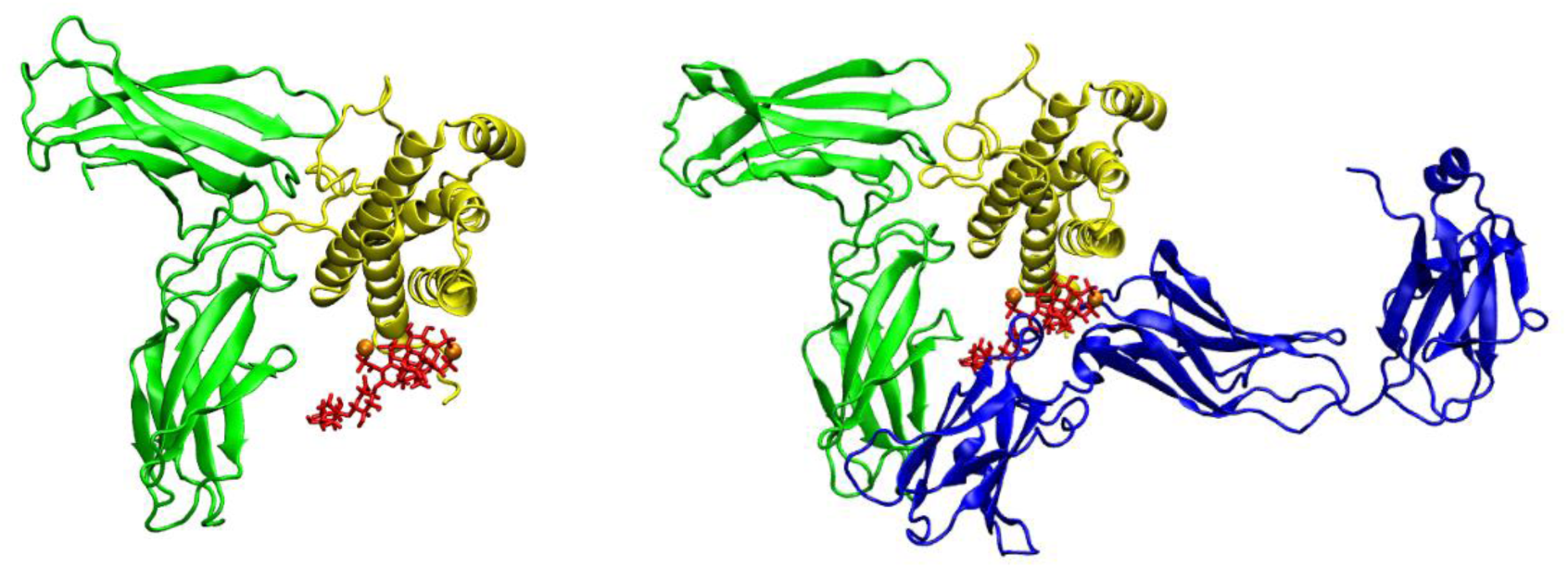
The IL6/IL-6Rα/LMWH/Mg complex. (A) IL-6 in yellow, heparin in red, IL-6Rα in green; Mg^2+^ ions presented as orange spheres; (B) The same complex, with gp130 receptor (in blue) superimposed.

## 4 Discussion

The severe phase of COVID-19 is characterised by a high level of expression of interferon-stimulated genes with proinflammatory activities, showing the immunopathological role of IFNγ response [4]. Among the elevated cytokines, IFNγ and the IFNγ-stimulated chemokines (such as IP-10 (CXCL10), MCP-1 (CCL2), and MIG) are especially prominent already in the very early fever days. The over-production of pro-inflammatory IFNγ and chemokines might be resulting from the lack of IL-10 – mediated down-regulation of the immune responses to SARS-CoV infection [54]. The elevated levels of IFNγ may also be related to lymphopenia since IFNγ was reported to induce apoptosis of activated T cells [55].

IFNγ can induce both pro- and anti-inflammatory responses. This allows it to tailor the immune response either for defense against infection or towards maintaining the homeostasis of the host [56]. When overexpressed, IFNγ is a key factor in the pathogenesis of autoimmune diseases, inflammation, atherosclerosis, neurodegenerative diseases, and cancer [56–58]. Besides CRS, the treatment of such diseases may also benefit from the inhibition of IFNγ signal transduction.

In fact, it is known that IFNγ binds with high affinity to glycosaminoglycans (GAGs), in particular heparan sulfate (HS) and heparin [59–61]. These interactions affect the activity, physico-chemical properties and proteolytic processing of the C-terminal domain of the cytokine. However, thus far the use of heparin as an effective IFNγ inhibitor directly preventing IFNγ from binding to its extracellular receptor was given little to no attention.

Our experimental and computational investigations show that LMWH binds with high affinity to IFNγ and at a certain concentration is able to inhibit fully the interaction with its cellular receptor, thus blocking the IFNγ signalling pathway and the expression of the IFNγ induced proteins. This observation reveals heparin as a potent candidate for regulation of the IFNγ overexpression. In particular, it can be helpful in the therapy of autoimmune diseases, acute inflammatory processes, and cytokine release syndrome.

IL-6 signalling is achieved via two different mechanisms. The first one, the cis-signalling, requires binding of IL-6 to its membrane-bound IL-6 receptor. This complex subsequently recruits two molecules of membrane-bound gp130, a process that leads to downstream signalling via JAK1/STAT3 pathway kinases (which is very similar to the interferon signalling pathway) [62]. The second mechanism, the trans-signalling, depends on the binding of IL-6 to the soluble IL-6 receptor (sIL-6R), which is expressed via mRNA splicing or proteolysis [61]. The formation of IL-6/sIL-6R complex activates gp130 on the cells that do not express the membrane-bound IL-6 receptor [63]. The trans-signalling is responsible for most of IL-6 inflammation-inducing capabilities (including the cytokine storm), whereas the anti-inflammatory activities of the cytokine are mediated via the cis-signalling [64].

Levels of IL-6 are very low under normal conditions, but they can raise many thousand-fold in inflammatory states. This is the reason for IL-6 being used, similar to the C-reactive protein (CRP), as a biomarker for the inflammation levels in patients with cancer, infection, autoimmune diseases or COVID-19 [65,66].

Inhibition of the IL-6/gp130/STAT3 pathway is considered to have therapeutic potential in cancer, autoimmune diseases, infection and COVID-19 [67,68]. At present, an antibody that blocks site I – siltuximab, has been approved as a drug. Two other antibodies are in different trial phases. Neutralising antibodies that block IL-6 receptors outside of the cell or small chemicals that target the involved kinases and transcription factors within the cell have been developed, some are approved as biologics (such as tocilizumab) and others are in different trial stages. However, the major concern for their use is in side effects, increased risk of bacterial and viral infections or their lack of specificity [69].

One feasible alternative for blocking the binding of IL-6 to its receptors is the application of LMWH. Our simulations show that IL-6 interacts with heparin forming a stable complex. The heparin molecule blocks the binding site I of IL-6 and prevents helices A and D from getting in close contact with IL-6Rα. Thus heparin inhibits formation of the IL-6/IL-6Rα complex and breaks the IL-6 signaling path. The other way in which heparin influences the biological activity of IL-6 is through binding to the complex IL-6/IL-6Rα blocking the binding site II of IL-6 and preventing helices A and C to be in a proper position to bind to gp130 receptor. These findings explain experimental observations [67] on heparin binding properties of IL-6. All together brings us to the conclusion that LMWH is a strong candidate for IL-6 activity inhibitor especially in the context of trans-signalling mechanism.

GAGs are universally present on cell surfaces and might be expected to serve as a broad spectrum and relatively non-specific receptors for virus binding [70]. Heparin as a potential inhibitor of viral attachment has been demonstrated for DENV [71]. Recently, viral attachment to GAGs was reported for IFNα/βBP from variola (smallpox virus), monkeypox viruses [72], hepatitis B [73], and Japanese encephalitis virus (JEV) [74]. Heparin and its derivatives have been shown to be effective in preventing the infection of cells by the influenza virus, strain H5N1 [75], and to have effects on ZIKV-induced cell death that are independent of adhesion and invasion [76]. Recently it was shown that heparin binds to the SARS-CoV-2 Spike protein receptor binding domain and induces conformational changes in it. As a result, heparin inhibits virus invasion by 70% [77].

GAGs and heparin in particular have been suggested as playing a protective role in the inflammation response [78], which has been interpreted as involving (among other things) the inhibition of elastase and the interaction with several cytokines [21]. Heparin and its analogues have been proposed for the application to elastase inhibition for cystic fibrosis treatment (or other conditions, such as acute respiratory distress syndrome (ARDS). Many of the applications of heparin may stem, at least in part, from the ability of heparin to moderate inflammation (see [19,79]). In general, heparin is perceived as acting via several routes, among them interaction with cytokines [80].

In the case of COVID-19 the heparin is used as an anticoagulant drug following the observation that patients in acute stage develop diffuse alveolar damage with extensive pulmonary coagulation activation resulting in fibrin deposition in the microvasculature and formation of hyaline membranes in the air sacs [81]. A wide multi-centre international trial on treatment with nebulised heparin of patients with COVID-19 was initiated [82]. COVID-19 patient autopsies have revealed thrombi in the microvasculature, suggesting intravascular coagulation as a prominent feature of organ failure in these patients. Elevated D-dimer levels were identified as a clear indicator for severity and mortality. Reduction of the hospital stay, decrease of severity and reduction of the mortality was registered in patients treated with heparin [83]. In another study with 449 patients involved and 99 out of them treated mainly with LMWH, the D-dimer and prothrombin time were positively and the platelet counts were negatively correlated with mortality. A reduced mortality in severe patients treated with heparin was reported in [84]. A similar response – reduced severity and mortality – was observed also in hemodialysis patients who were treated with heparin or other anticoagulants according to the standard protocols [85].

The dynamic changes of lymphocyte subsets and cytokines profiles of patients with COVID-19 and their correlations were studied in [86]. The severe cases showed a significant sustainable decrease in lymphocyte counts and an increase in neutrophil one compared to the respective levels in mild cases. In particular, decrease in the levels of CD4+ and CD8+ T cells and increase of IL-2, IL-6, IL-10 and IFNγ were registered. Similar results were reported in [87] indicating an overall decline of lymphocytes including CD4+ and CD8+ T cells, B cells, and NK cells in severe patients. The number of immunosuppressive regulatory T cells was increased in mild patients. A remarkable up-regulation of inflammatory cytokines IL-2, IL-6, and IL-10 was registered in severe patients. It was shown that the comprehensive reduction of the lymphocyte levels, and the elevation of IL-2 and IL-6 are reliable indicators of severe COVID-19. In a retrospective clinical study [88] it was reported that LMWH not only improves the coagulation dysfunction of COVID-19 patients but also significantly reduces the levels of IL-6 and increases levels of the lymphocytes.

In summary, the clinical studies show a significant increase of the levels of IL-2, IL-6, IL-10, and hINFγ at the beginning of the acute phase of the disease. At the same time, a reduction of the levels of CD4+ and CD8+ T-cells and B-cells is observed. After patient treatment with LMWH, a fast reduction of the inflammatory cytokines, restoration of the levels of CD4+ and CD8+ T-cells and B-cells, reduction of the levels of virions and significantly faster recovery was observed. Combined with the clinical studies, our results suggest that heparin, in particular LMWH, is a very promising candidate for a therapeutic means for prevention and inhibition of the cytokine storm in COVID-19 patients.

## 5 Conclusions

There is a considerable volume of observational data regarding the use of heparin in the treatment of COVID-19. The reason for including this drug in the treatment protocols is to reduce the formation of blood clots that prevents a number of complications (heart attacks, emboli, strokes, etc.), but also obstruction of the capillaries in the alveoli.

The main result of our study is that heparin is, in fact, a potent anti-inflammatory agent that can be used in acute inflammatory conditions, due to its ability to block at least two (IFNγ and IL-6) of the key signaling molecules involved in the process of the inflammation development. Thus, it should be prescribed in the initial stages of an acute condition (pneumonia). The decision to start application of LMWH in COVID-19 patients and its dosage should be based on careful follow up during the treatment course of the levels of D-dimer, prothrombin time, platelet counts, IL-2, IL-6, IL-10, IFNγ, IP-10 (CXCL10), MCP-1 (CCL2), MIG, CD4+ and CD8+ T-cells and B-cells. Elevation of D-dimer, IL-2, IL-6, IFNγ (or IP-10) levels, along with a reduction of CD4+ and CD8+ T-cells and B-cells counts is a clear signal for the approaching onset of the acute phase of the disease and an indication for the inclusion of LMWH in the treatment plan. The doses still need to be elaborated but our results suggest an orientation towards the upper limit of the dosage scale, whereby the action of heparin will be threefold – anticoagulant, anti-inflammatory and antiviral.

The action of heparin as an anti-inflammatory agent does not depend on the virus type and, in general, the cause for the acute inflammatory process. This broadens its application range even more – for treatment of viral infections leading to acute inflammatory conditions characterised by increased cytokine levels, in particular those of IL-6 and IFNγ.

Heparin is a well-known and widely used medication and its side effects are studied in detail. This greatly facilitates the expansion of its scope towards the treatment of acute inflammatory conditions, in particular those associated with COVID-19; the general limitations for LMWH use remain valid in these cases as well.

Overexpression of IFNγ is also observed in a number of autoimmune diseases. A possible treatment strategy consists in inhibiting the biological activity of IFNγ. The results of our study reveal the capability of heparin to influence the development of such autoimmune diseases.

Our findings about the potential of heparin as an anti-inflammatory agent need validation through appropriate clinical trials. In view of the current situation with the COVID-19 pandemic and the extensive use of heparin in the clinical practice as anticoagulant drug, we strongly suggest the wide application of heparin for COVID-19 treatment as a potent anti-inflammatory ligand in the framework of carefully monitored and controlled clinical trials.

## SUPPLEMENTARY MATERIAL

**Fig. S1.**
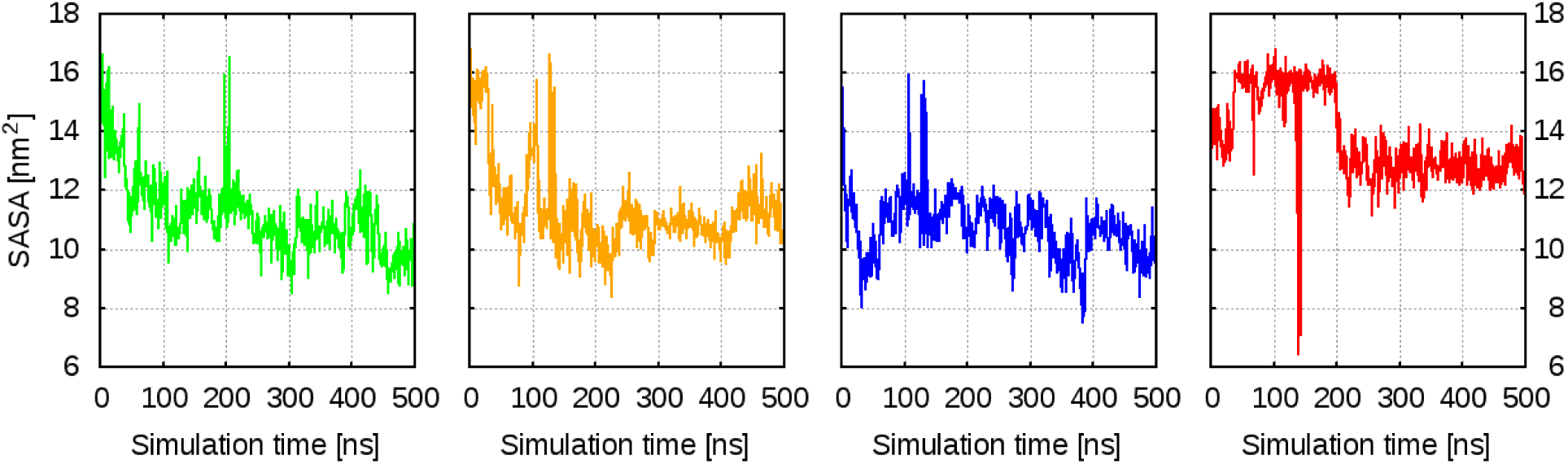
Solvent accessible surface area (SASA) of the hexasaccharides: dp6_1 – green curve, dp6_2 – orange curve, dp6_3 – blue curve, and dp6_4 – red curve.

**Fig. S2.**
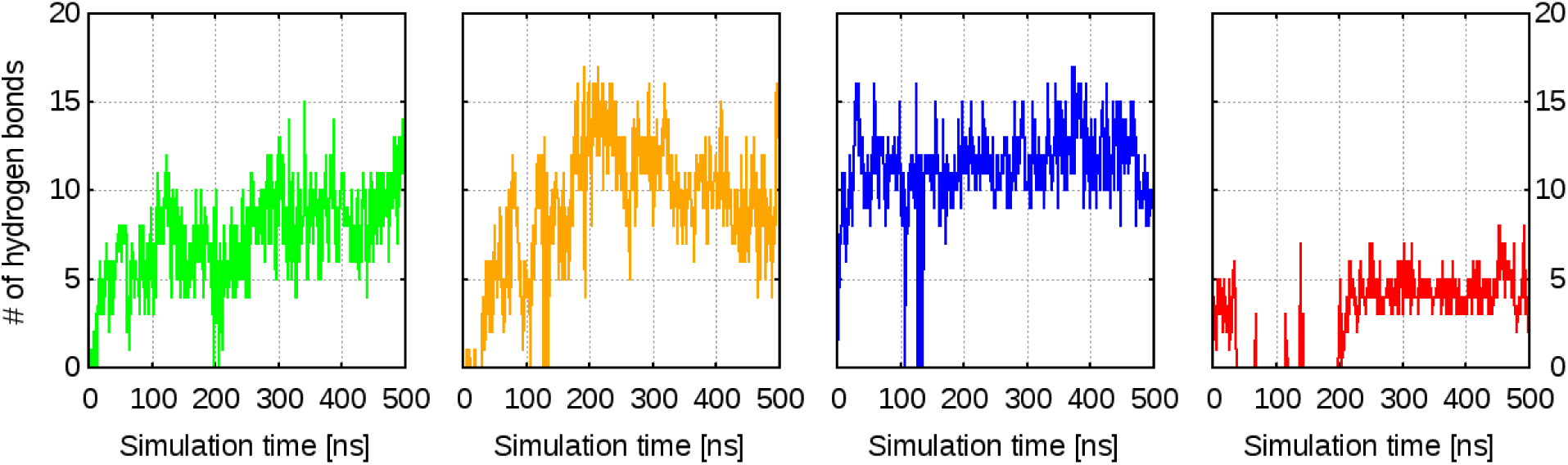
Hydrogen bonds formed between hIFNγ and each of the hexasaccharides: dp6_1 – green curve, dp6_2 – orange curve, dp6_3 – blue curve, and dp6_4 – red curve.

**Fig. S3.**
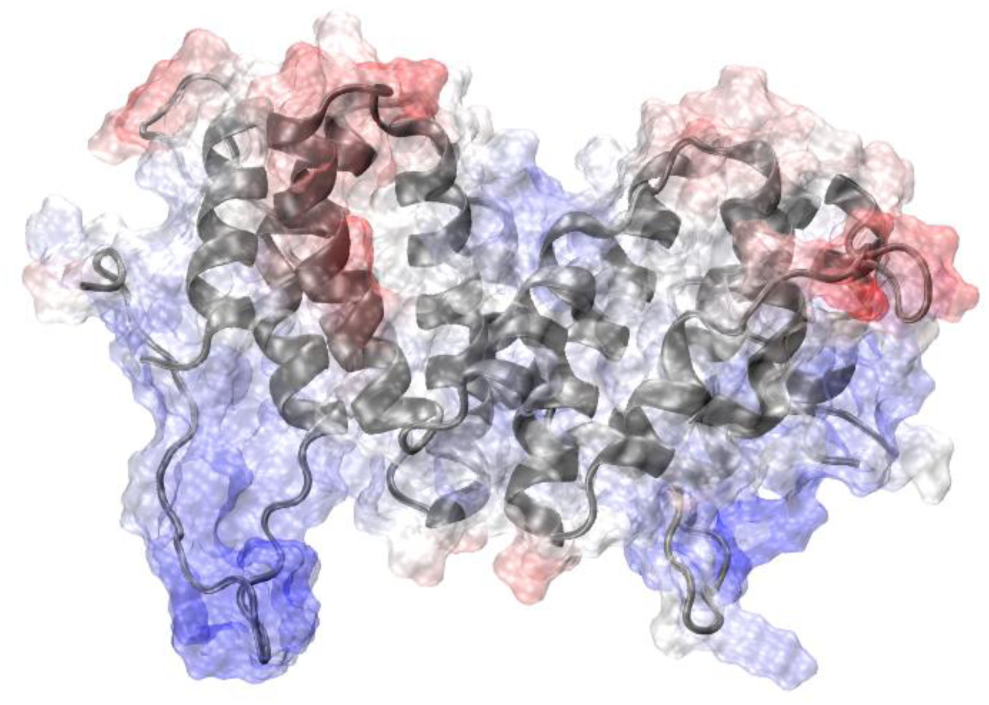
Electrostatic potential surface of the starting structure of hIFNγ. Positively charged areas are coloured in blue, negatively charged areas – in red.

## Authors’ contributions

L.L. designed and coordinated the study, wrote the initial draft, and did the final editing; G.N. designed the experimental studies, conducted together with E.K., K.M., A.G., and R.H; P.P., M.R., E.L., and N.T. performed the MD simulations; P.P., M.R., E.L., and E.K. contributed texts to the initial draft, G.N. and N.I. contributed substantially; the whole team participated in data analysis and interpretation; G.N. and A.G. edited the manuscript; N.I. performed extensive editing; G.N. and N.I. coordinated the work, arranging also for laboratory- and computational resources respectively.

## Declaration of interests

The authors declare no conflict of interests. The funders had no role in the design of the study; in the collection, analyses, or interpretation of data; in the writing of the manuscript, or in the decision to publish the results.

## Acknowledgements

This work was supported in part by the Bulgarian National Science Fund under Grant DN-11/20/2017 and by the Bulgarian Ministry of Education and Science, under the National Research Programme “ICTinSES” (Grant D01-205/23.11.2018, approved by DCM # 577/17.08.2018). EK acknowledges the support from the National Research Program “Young scientists and postdoctoral students” of the Bulgarian Ministry of Education and Science (DCM # 577/17.08.2018). Computational resources were provided by the Centre for Advanced Computing and Data Processing, supported under Grant BG05M2OP001-1.001-0003 by the Science and Education for Smart Growth Operational Program (2014-2020) and co-financed by the European Union through the European structural and investment funds. We acknowledge the constructive comments from Dr. A. Pashov, PhD, of the Institute of Microbiology at BAS.

## Notes

### Competing Interest Statement

The authors have declared no competing interest.

